# CRISPR-guided programmable self-assembly of artificial virus-like nucleocapsids

**DOI:** 10.1101/2020.10.17.343996

**Authors:** Carlos Calcines-Cruz, Ilya J. Finkelstein, Armando Hernandez-Garcia

## Abstract

Designer virus-inspired proteins drive the manufacturing of more effective and safer gene-delivery systems as well as simpler models to study viral assembly. However, the self-assembly of engineered viromimetic proteins on specific nucleic acid templates, a distinctive viral property, has proved difficult. Inspired by viral packaging signals, we harness the programmability of CRISPR-Cas12a to direct the nucleation and growth of a self-assembling synthetic polypeptide into virus-like particles (VLP) on specific DNA molecules. Positioning up to ten nuclease-dead Cas12a (dCas12a) proteins along a 48.5 kbp DNA template triggers particle growth and full DNA encapsidation at limiting polypeptide concentrations. Particle growth rate was further increased when dCas12a was dimerized with a polymerization silk-like domain. Such improved self-assembly efficiency allows for discrimination between cognate versus non-cognate DNA templates by the synthetic polypeptide. Our CRISPR-guided VLPs could help develop programmable bio-inspired nanomaterials with applications in biotechnology as well as viromimetic scaffolds to improve our understanding of viral self-assembly.

## Introduction

Virus-like particles (VLPs) mimic the capability of some viruses to encapsulate and protect genetic material from degradation by nucleases. We have previously described VLPs formed by the self-assembly of a tri-block polypeptide (C-S_10_-B) that functionally mimics the tobacco mosaic virus coat protein^1^ (Fig. 1a). C-S_10_-B fuses three independent blocks: 1) “C” (∼400 aa), a random coil collagen-like domain that consists mostly of glycine, proline and uncharged polar amino acids^2^; 2) “S_10_”, a silk-inspired polymerization domain with the sequence [(AG)_3_QG]_10_ that is responsible for C-S_10_-B self-assembly into rod-like structures^3–5^; and 3) “B”, a cationic dodecalysine stretch that interacts electrostatically with nucleic acids and other polyanions^6, 7^. C-S_10_-B nucleates (without sequence specificity) on double stranded DNA (dsDNA), albeit with a preference for free DNA ends^8^. After rate-limiting nucleation, C-S_10_-B filaments grow rapidly through elongation^1, 9^, similar to the assembly of the tobacco mosaic virus coat protein on the genomic ssRNA.

**Fig. 1:**
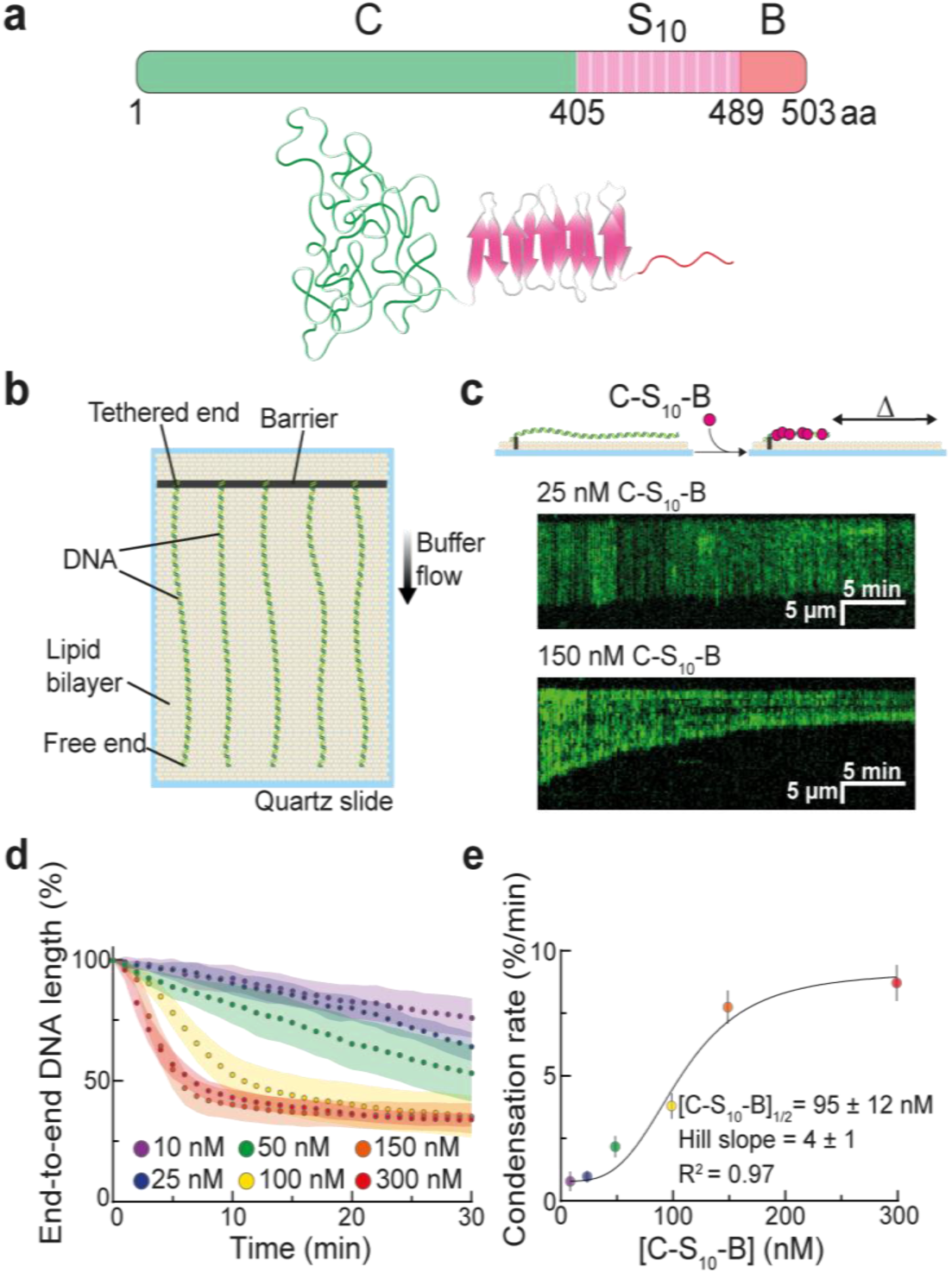
The synthetic polypeptide C-S_10_-B packages DNA into nucleocapsids via molecular self-assembly. **a**: Schematic depiction and modular design of C-S_10_-B. **b**: Illustration of the DNA curtain assay. Individual DNA molecules are captured on the surface of a lipid-coated flowcell and extended by gentle buffer flow. **c**: Representative kymographs showing condensation of individual DNA strands (green) at different rates determined by C-S_10_-B concentration. The DNA is stained with the intercalating dye YOYO-1. **d**: DNA condensation profiles at the indicated C-S_10_-B concentrations. The circles and shaded areas represent the mean and standard deviation for 25 molecules per condition, respectively. **e**: DNA condensation rate in the first 3 min of assembly at different concentrations of C-S_10_-B. The data were fitted to a sigmoid (solid line) and shown are mean ± 95% confidence intervals for 25 DNA molecules per condition.

Viral coat proteins preferentially encapsulate their own genomes. They achieve such specificity by encoding one or more packaging signals along the viral DNA or RNA. These sequences bind capsid proteins with high affinity^10–17^ and decrease the energy barrier for nucleation, thereby promoting encapsidation of the viral genome among a vast excess of cellular nucleic acids^18–21^. We reasoned that designer VLPs can also leverage a similar packaging signal to enhance nucleation at specific DNA sites. We chose the catalytically dead CRISPR-Cas12a (dCas12a) as a programmable nucleation signal because it binds dsDNA with 50 fM affinity, has a higher DNA binding specificity than *S. pyogenes* Cas9^22^, and can be directed to multiple sites along the DNA via pooled CRISPR RNAs (crRNAs)^23^.

We decorated a DNA template with multiple dCas12a or dCas12a coupled to the S_10_ polymerization domain and favored DNA encapsidation by the synthetic polypeptide (C-S_10_-B). Self-assembly kinetics were monitored in real-time using the single-molecule DNA curtain assay (Fig. 1b)^24, 25^. For this, arrays of DNA molecules (48.5 kbp, derived from λ-phage) are affixed to a lipid bilayer via a biotin-streptavidin linkage in a microfluidic flowcell. Microfabricated chrome barriers are used to organize thousands of DNA strands for high-throughput data collection and analysis. The surface-immobilized DNA is extended for fluorescent imaging via the application of mild buffer flow. DNA length was monitored along the experiment since time-dependent DNA contraction is a direct readout of VLP filamentation (Fig. 1c)^1^.

## Results

First, we identified the minimal concentrations for efficient encapsidation of individual dsDNA substrates by C-S_10_-B alone. At 100-300 nM, C-S_10_-B monotonically condensed individual DNA molecules until complete assembly of linear particles with a final length of 34 ± 3% of the initial length (N = 25 DNA molecules at each concentration) (Fig. 1d). Total DNA contraction to about one third of the DNA original length has also been observed by previous AFM studies^1^. Concentrations between 10 nM and 50 nM decreased the DNA encapsidation rate (measured as t_half_, the time required to achieve half of total packaging) and did not completely contract the DNA to its final encapsulated length, suggesting limited C-S_10_-B binding. Initial condensation rate increased with polypeptide concentration in a sigmoid-shaped curve distinctive of nucleated self-assemblies (Fig. 1e). Nucleation and particle growth were significantly impaired at 25 nM C-S_10_-B compared with 150 nM (Fig. S2). These results are consistent with a dynamic equilibrium between C-S_10_-B nucleation-filamentation and dissociation from DNA, akin to RAD51 and other dynamic filaments^26^.

Next, we evaluated whether positioning dCas12a on the DNA can improve encapsidation at low C-S_10_-B concentrations. The DNA was uniformly decorated with five or ten dCas12a-crRNA ribonucleoproteins (RNPs) (Fig. 2a). We confirmed site-specific target binding by imaging fluorescent RNPs along the DNA molecule (Fig. S3a). We observed from 1 to 4 (1.2 ± 0.5 RNP per DNA) target-bound RNPs on DNA substrates harboring five binding sites (N = 456 DNA molecules) and 1 to 7 RNPs (2.6 ± 1.1 RNP per DNA) on DNA substrates with ten binding sites (N = 185 DNA molecules) (Fig. S3b). Because not all RNPs are decorated with fluorescent QDs, our results are a lower bound on the true target site occupancy. To monitor how encapsidation varies with RNP density, we injected 25 nM C-S_10_-B into curtains with DNA strands pre-decorated with either five or ten RNPs (Figs. 2b-c). DNA contraction rate increased two-and three-fold for DNA decorated with five (t_half_ = 15 ± 7 min; N = 25 DNA molecules) or ten (t_half_ = 10 ± 3 min; N = 25) RNPs with respect to naked DNA (t_half_ = 31 ± 6 min; N = 25) (Fig. 2d). We conclude that dCas12a RNPs can be installed at specific sites along the DNA template and that such decoration promotes and accelerates encapsidation by C-S_10_-B.

**Fig. 2:**
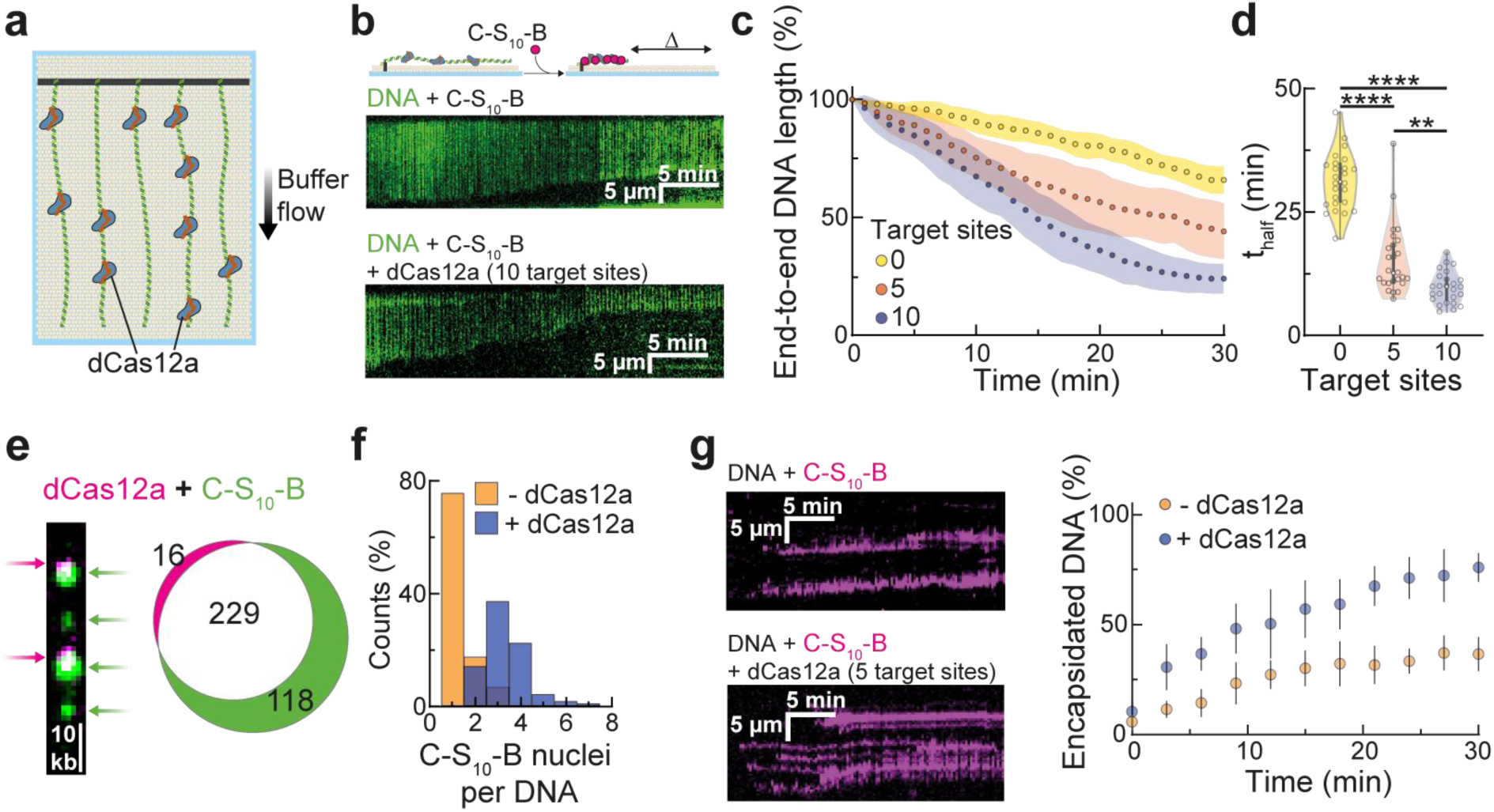
Seeding assembly with dCas12a improves DNA encapsidation by C-S_10_-B. **a**: Illustration of a dCas12a-decorated DNA substrates. dCas12a is incubated with pools of crRNAs and directed to 5 or 10 sites uniformly distributed along the DNA prior to C-S_10_-B injection. **b**: Representative kymographs show faster encapsidation at 25 nM C-S_10_-B after the DNA (green) is decorated with dCas12a. **c**: Condensation profiles at 25 nM C-S_10_-B for naked DNA, and DNA decorated with dCas12a targeting 5 or 10 sequences along the template. The circles and shaded areas represent the mean and standard deviation for 25 molecules per condition, respectively. **d**: Violin plots showing the time (t_half_) required to reach half of maximum condensation for each DNA molecule analyzed in (c). For zero target sites, we fitted the curve to a sigmoid and extrapolated t_half_ for molecules that did not reach 65.5% encapsidation during the experiment (30 min). (**) and (****) indicate p < 0.01 and p < 0.0001, respectively. **e**: Fluorescent dCas12a (magenta, 93%) co-localizes with fluorescent C-S_10_-B clusters (green, 66%). **f**: Positioning five dCas12a molecules on the DNA increases the number of fluorescent C-S_10_-B clusters relative to undecorated DNA (N = 484 and 246 clusters, respectively). **g**: Kymographs showing C-S_10_-B binding (magenta) on undecorated (top) or dCas12a-decorated (bottom) DNA and percentage of the DNA strand length that is encapsidated by the fluorescent C-S_10_-B. Shown are the mean standard deviation for 10 molecules per condition.

We reasoned that dCas12a organizes large C-S_10_-B clusters that further polymerize into filaments. Two-color imaging confirmed that fluorescent C-S_10_-B co-localizes with the RNPs at early stages of encapsidation (Fig. 2e). Positioning five RNPs on the DNA increased the number of fluorescent C-S_10_-B puncta per DNA strand from 1-3 (1.3 ± 0.6, N = 166 DNA molecules) to 2-7 (3.3 ± 1.0, N = 97 DNA molecules) for naked and dCas12a-decorated DNA, respectively (Fig. 2f). C-S_10_-B protomers can freely diffuse on the DNA^27^. In the presence of buffer flow, these molecules slide and assemble into large clusters at the free DNA ends. In contrast, C-S_10_-B accumulated at dCas12a sites on the decorated DNA (Fig. S3c). Filament growth was also more rapid on pre-decorated DNA substrates (t_half_ = 15 ± 6 min; N = 10 DNA molecules) than on naked DNA (t_half_ = 45 ± 15 min; N = 10 DNA molecules) (Fig. 2g). C-S_10_-B filamentation was unable to displace dCas12a, which forms a stable RNA:DNA loop (R-loop) with the DNA substrate (Fig. S3d). Because dCas12a may act as a roadblock for C-S_10_-B linear diffusion on the DNA, we also tested whether other DNA-binding proteins will also accumulate C-S_10_-B clusters. Notably, DNA decoration with nucleosomes also accelerated DNA encapsidation and particle growth (Fig. S4). We conclude that dCas12a stalls C-S_10_-B sliding on DNA, which enhances polypeptide collisions in its vicinity and triggers particle nucleation. In support of this model, Marchetti et al.^27^ observed that immobile C-S_10_-B clusters, and not their sliding counterparts, initiate filament growth.

To further promote and accelerate DNA packaging at limiting C-S_10_-B concentrations, we physically coupled dCas12a to the di-block polypeptide C-S_10_ via the rapamycin-inducible FKBP and FRB dimerization domains (Fig. 3a). We cloned and purified a dCas12a(FKBP) C-terminal fusion and verified that this construct retains target-specific DNA binding (Fig. S1). The FRB domain was fused to a truncated C-S_10_ polypeptide that lacks the “B” DNA-binding module. Both proteins, dCas12a(FKBP) and C-S_10_(frb), were incubated with rapamycin and the dimerized complex was injected into the flowcell prior to incubation with 10 nM C-S_10_-B (Fig. 3b-c, Fig. S5). Positioning five dimerized dCas12a-S10 complexes along the DNA substrate accelerated encapsidation by C-S_10_-B two-fold relative to dCas12a(FKBP) RNP alone (t_half_ = 9 ± 3 min; N = 25 vs. 18 ± 6 min; N = 25), and five-fold relative to the undecorated DNA (t_half_ = 46 ± 13 min; N = 25) (mean ± SD, Fig. 3d-e). We confirmed this result via ensemble biochemical electrophoretic mobility shift assays (EMSAs) (Fig. S6). Positioning the dCas12a-S10 complex on a 2.5 kbp linear dsDNA or a 9.5 kbp plasmid dsDNA also improved encapsidation at limiting C-S_10_-B concentrations (Fig. S6). These results indicate that initiating nucleation via interspersed dCas12a RNPs fused to the self-assembly domain S_10_ accelerates DNA packaging at sub-saturating C-S_10_-B concentrations.

**Fig. 3.**
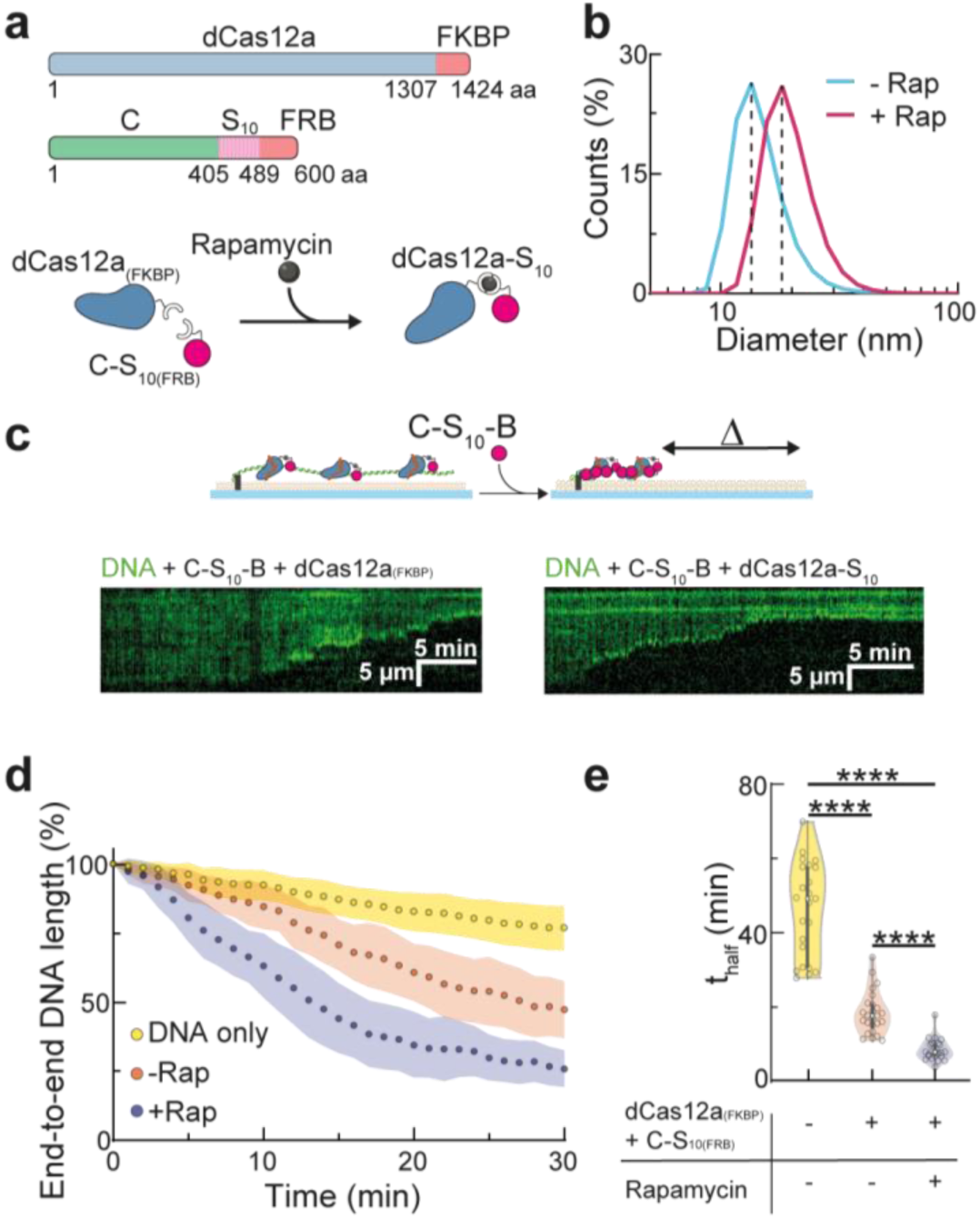
Coupling of dCas12a to the self-assembly domain C-S_10_ improves C-S_10_-B nucleation and assembly. **a**: Schematic depiction of the modular design of dCas12a_(FKBP)_ and C-S_10(frb)_ and their dimerization via rapamycin to form the dCas12a-S_10_ complex. **b**: Dynamic light scattering experiment showing rapamycin (Rap) induced dimerization of dCas12a_(FKBP)_ and C-S_10(frb)_. The shift in population size occurred immediately after rapamycin addition. **c**: Representative kymographs showing that decorating DNA with dCas12a-S_10_ accelerates DNA packaging at 10 nM C-S_10_-B relative to decoration with dCas12a_(FKBP)_ alone. **d**: Condensation profiles at 10 nM C-S_10_-B for undecorated DNA, and DNA decorated with 5 dCas12a_(FKBP)_ or 5 dCas12a-S_10_. The circles and shaded areas represent the mean and standard deviation for 25 molecules per condition, respectively. **e**: Violin plots showing the time (t_half_) required to reach half of maximum condensation for each DNA strand analyzed in (d). Extrapolation (after sigmoid curve fitting) was used to estimate t_half_ for molecules that did not reach 65.5% encapsidation during the experiment (30 min). (****) indicates p < 0.0001.

Finally, we assessed whether targeting dCas12a-S10 to a specific DNA template can trigger VLP encapsidation in the presence of other (non­cognate) DNA molecules. For this assay, we immobilized an equimolar mixture of two DNA templates on the flowcell surface; one was 20 kbp and the second was 28.5 kbp (Fig. 4a). dCas12a-S10 was directed to 5, 10 or 25 sites on the 28.5 kbp template (Fig. 4b). As seen in Fig. 4c-d and Fig. S7, dCas12a-S10 selectively accelerated the packaging of the target DNA template and the encapsidation rate increased three-fold from zero target sites (t_half_ = 62 ± 26 min; N = 25) to a maximum of 25 target sites (t_half_ = 19 ± 6 min; N = 25).

**Fig. 4.**
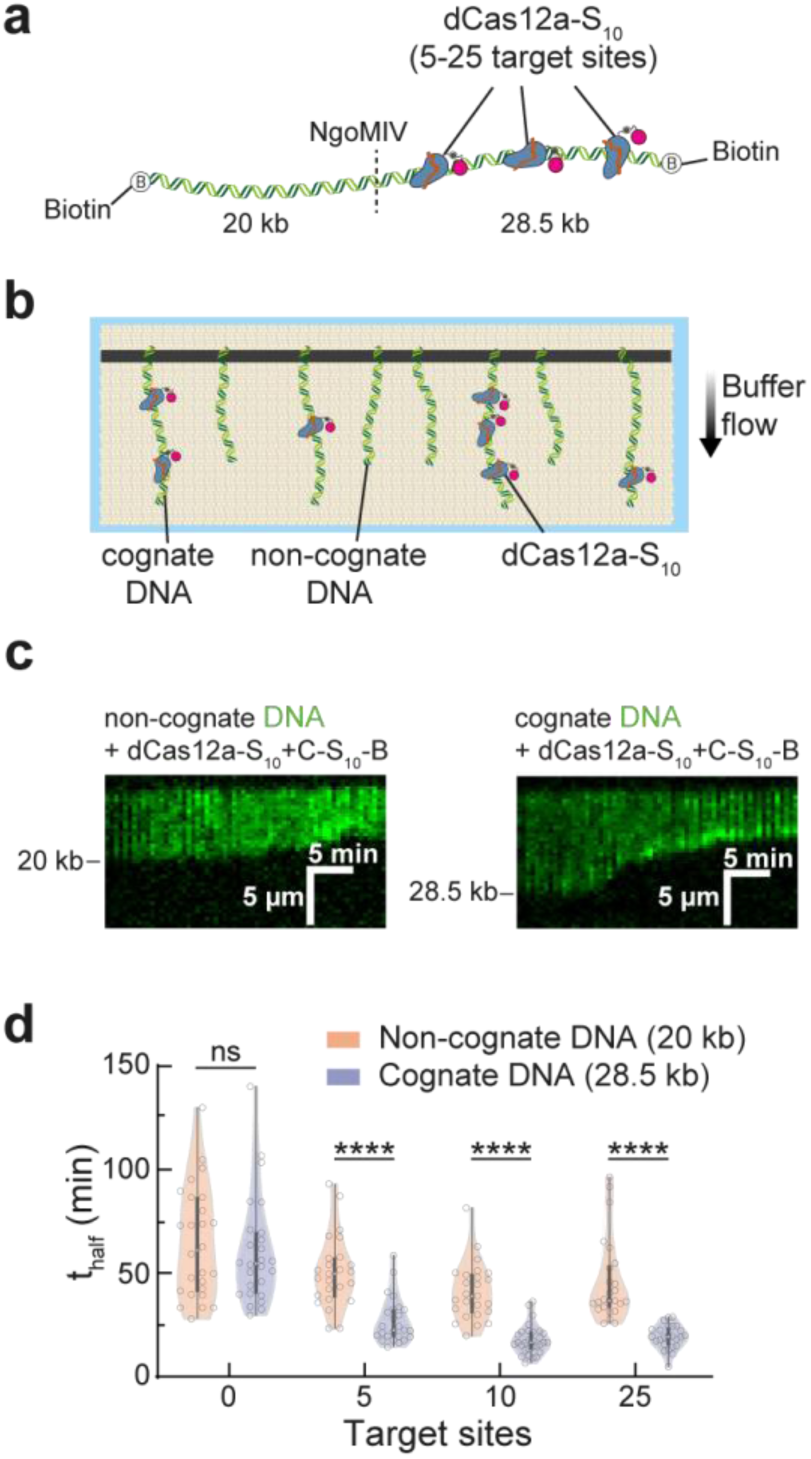
C-S_10_-B selectively assembles on DNA that is decorated with dCas12a-S_10_. **a**: DNA was ligated with biotinylated oligos at both ends followed by cleavage with NgoMIV to generate two biotinylated DNA substrates (20 kbp and 28.5 kbp) distinguishable by size after YOYO-1 staining. **b**: The 28 kbp DNA (cognate DNA) was decorated with 5-25 dCas12a-S_10_ prior to incubation with 10 nM C-S_10_-B. **c**: Representative kymographs showing encapsidation of the non­cognate (20 kbp) and cognate (28.5 kbp) DNA after decoration with 10 dCas12a-S_10_. **d**: Violin plots showing the time (t_half_) required to reach half of maximum condensation for cognate and non-cognate DNA strands after decoration with 0-25 dCas12a-S_10_. We estimate (after sigmoid curve fitting) t_half_ for molecules that did not reach 65.5% encapsidation during the experiment (30 min). (****) indicates p < 0.0001.

## Discussion

Taken together, our data show that a target DNA-bound dCas12a (or any strongly DNA-bound roadblock protein) can serve as a viral-like packaging signal during the self-assembly of VLPs—especially if the roadblock protein contains a self-assembly or polymerization domain. RNA-guided CRISPR-Cas proteins are especially attractive as artificial packaging signals because they can be targeted to any DNA sequence that is proximal to a protospacer adjacent motif (PAM). For AsdCas12a, the PAM consensus sequence is TTTV; this PAM appears on average every 32 bp on 48.5 kbp long λ-phage DNA. Other CRISPR-Cas enzymes, including Cas9, can also serve as nucleation signals. Relaxed PAM variants, both for Cas9 and Cas12, will further increase the targeting possibilities for rapid and sequence specific VLP assembly^28–31^. Additionally, this work shows that C-S_10_-B protein can be re­engineered to more closely resemble virus-like features.

In summary, we have used CRISPR-dCas12a protein and coupled it to the polymerization domain S_10_ to form a “packaging-signal recognition complex” (in analogy with the assembly of viral capsid proteins around the viral genome) that triggers binding of C-S_10_-B and packaging of target DNA sequences. Importantly, such packaging recognition complex (and hence artificial particle nucleation and growth) can be easily redirected toward different DNA sequences by simple design of the crRNA without need of the more cumbersome manipulation of protein domains. Moreover, our results highlight the importance of having multiple strong and specific interactions in templated nucleoprotein self-assemblies and should prove useful for developing new tailor-made nanomaterials for diverse biotechnological applications.

## Materials and Methods

### Plasmids, protein expression and purification

The FK506 Binding Protein (FKBP) and FKBP-Rapamycin Binding domain (FRB) sequences were extracted from pGEX-2T plasmids (kindly provided by Tom Wandless, Stanford University) and inserted at the C-terminus of nuclease inactivated version of AsCas12a (D908A) and C-S_10_, respectively.

The pET19 plasmids with dCas12a and dCas12a_(FKBP)_ (both with an N-terminus 6XHis-SUMO tag) were used to transform BL21 (DE3) *Escherichia coli.* Colonies were grown on Luria Bertani plates with 100 ug/mL ampicillin and used to inoculate Terrific Broth liquid culture in Fernbach flasks. Cultures were grown at 37°C until OD_6_oo ∼0.8 and then transferred to 12°C for 1 h before induction with 1 mM IPTG. Cultures were then grown for 24 h at 12°C and the pellets were collected by centrifugation and stored at −70°C.

The pellets were resuspended in buffer A (20 mM Tris-HCl pH 8.0, 250 mM NaCl, 10 mM imidazole) supplemented with 1 mM PMSF and sonicated on ice. The lysate was clarified by centrifugation at 30 000 g for 45 min and the supernatant containing the protein of interest was filtered through 0.2 μm syringe filters. The supernatant was then loaded into a 5 mL HisTrap (GE Healthcare) column, washed with buffer B (20 mM Tris-HCl pH 8.0, 1 M NaCl, 25 mM imidazole) and eluted with buffer C (20 mM Tris-HCl pH 8.0, 1 M NaCl, 250 mM imidazole). The eluted fraction was loaded into a dialysis membrane with 2.4 μm SUMO protease and dialyzed overnight at 4°C in buffer D for dCas12a (50 mM phosphate buffer pH 6.0, 100 mM KCl, 5 mM MgCh, 10% glycerol, 2 mM DTT) and buffer E for dCas12a_(FKBP)_ (20 mM HEPES-KOH pH 7.2, 100 mM KCl, 5 mM MgCl_2_, 10% glycerol, 2 mM DTT). The 0.2 μm filtered cleaved product was injected in a 5 mL HiTrap SP HP (GE Healthcare) cationic exchange column and eluted with a gradient to buffer F (20 mM HEPES-KOH pH 7.5, 2 M KCl). The fractions containing the protein of interest were pooled, concentrated with 100 kDa Amicon centrifugal filter units and injected into a size exclusion HiLoad 16/600 Superdex 200 pg column (GE Healthcare) equilibrated with storage buffer G (20 mM HEPES-KOH pH 7.5, 500 mM KCl, 10% glycerol). Purified proteins were concentrated to 10-12 μΜ, flash frozen in liquid nitrogen and stored at −70°C until use. Protein identity and purity were assessed with western blots and SDS-PAGE, respectively.

pPIC9 plasmids with C-S_10_-B and C-S_10(FRB)_ were linearized with SacI and electroporated into histidine auxotrophic *Pichia pastoris* GS115. Mut_+_ colonies expressing the protein were used to inoculate 300 mL of MGY and grown for 24 h until Od_6_oo ∼6.0. Cells were pelleted and resuspended in MM medium for protein expression for 72 h, with 1% methanol addition every 24 h. Culture supernatant was collected by centrifugation, the medium pH was adjusted to 8.0 with NaOH and 1 mM PMSF and 5 mM EDTA were added to inhibit proteolysis followed by 0.2 μm filtration.

For C-S_10_-B, the culture supernatant was then saturated to 50% with ammonium sulfate and incubated overnight at 4°C. The protein precipitate was resuspended in Milli-Q water at 65°C and the precipitation step was repeated once. The precipitate was resuspended in Milli-Q water at 65°C before 50 mM NaCl and 40% acetone addition. The sample was centrifuged to discard precipitates and the concentration of acetone in the supernatant was increased to 80% to selectively precipitate C-S_10_-B. The precipitate was air dried for 15 min, resuspended and dialyzed overnight against Milli-Q water at 4°C, followed by flash freeze and lyophilization. Before use, the lyophilized protein was dissolved in Milli-Q water at 0.1 mg/mL and incubated at 70°C for 15 min. For C-S_10(FRB)_, the culture supernatant was concentrated with 30 kDa Amicon stirred cells and proteins were precipitated with 80% saturation ammonium sulfate at 4°C overnight. The protein precipitate was resuspended in PBS pH 7.5 and the precipitation step was repeated once. The precipitate was resuspended in PBS pH 7.5 and 40% acetone was added followed by centrifugation. The concentration of acetone in the supernatant was increased to 80% to selectively precipitate C-S_10(FRB)_. The precipitate was resuspended to ∼80 μΜ in PBS pH 7.5, dialyzed overnight against PBS pH 7.5 at 4°C and flash frozen. Protein identity and purity were assessed with mass spectrometry and SDS-PAGE, respectively.

### crRNA pools

The 24 nt forward oligo with the promoter for T7 RNA polymerase, and the 68 nt reverse oligos with the 24 nt sequence complementary to the forward oligo plus the 44 nt crRNA sequence were purchased from IDT. The T7 promoter forward oligo was mixed with pools of reverse oligos (1.5:1.0 molar ratio) and hybridized by incubation at 75°C for 5 min and then cooling to 25°C over 25 min in annealing buffer (10 mM Tris-HCl pH 7.5, 50 mM NaCl, 1 mM EDTA). The pool of partially double stranded DNA templates was used for *in vitro* transcription with HiScribe™ T7 Quick High Yield RNA Synthesis Kit (New England BioLabs) and the RNA was purified with TRizol (Ambion). crRNA integrity and purity were assessed with 15% acrylamide Urea-PAGE.

### Preparation of DNA substrates

To prepare the DNA template for curtain assays λDNA (125 μg, NEB) was incubated with 2 μΜ biotinylated oligo complementary to one of the two 12 nt cohesive ends in λDNA in T4 DNA ligase reaction buffer (NEB) and hybridized at 70°C for 15 min followed by cooling to 15°C over 2 h. T4 DNA ligase (2000 units, NEB) was added and the mixture was incubated at room temperature overnight. The ligase was inactivated with 2 M NaCl and the sample was injected into a Sephacryl S-1000 size exclusion column (GE Healthcare) to remove excess oligos and DNA ligase.

For experiments with nucleosomal DNA, purified biotinylated DNA was precipitated with sodium acetate (pH 5.5) and 0.3 M isopropanol to 1:1 v/v on ice, and centrifuged at 15 000 g for 30 min. The precipitate was resuspended to 0.8 nM and incubated with human histone octamers 3xHA H2A and H2B, H3, H4; Histone Source) at molar ratios of 50:1 and 00:1 (histone octamer to λDNA) in 2 Μ TE buffer (10 mM Tris-HCl pH 8, 1 mM EDTA, 2 Μ NaCl) with 1 mM DTT. The 100 μL mixture was loaded in dialysis buttons (10 kDa MWCO, BioRad) and dialyzed against dialysis buffer (10 mM Tri-HCl pH 7.6, 1 mM EDTA, 1 mM DTT and NaCl). The first 1 h dialysis step was done against 1.5 M NaCl, followed by 2 h dialysis steps against 1, 0.8, 0.6 and 0.4 mM with a final overnight step against 0.2 M NaCl at 4°C.

For experiments with two DNA strands, λDNA was ligated with biotinylated oligos complementary to both cohesive ends. Hybridization and ligation were performed as described above except that CutSmart buffer (NEB) supplemented with 1 mM rATP was used instead of T4 ligase buffer. The mixture was heated at 65°C for 10 min to inactivate the ligase and NgoMIV was added for DNA cleavage at 37°C for 3 h. Lastly, 1 M NaCl was added to the solution prior to size exclusion chromatography. Complete cleavage by NgoMIV was assessed with gel electrophoresis. All DNA substrates were stored at 4°C until use.

### Single-tethered DNA curtain assay

A solution containing 20 μL of a liposome stock solution (97.7% DOPC, 2.0% DOPE-mPEG2k and 0.3% DOPE-biotin) and 980 μL of lipid buffer 10 mM Tris-HCL pH 8, 100 mM NaCl) was injected into the assembled owcell and incubated for 30 min at room temperature. The flowcell was washed with BSA buffer (40 mM Tris-HCl pH 8, 2 mM MgCl_2_, 0.2 mg/mL BSA) and incubated for 10 min before injection of 0.1 mg/mL streptavidin in BSA buffer followed by 10 min incubation. The biotinylated λDNA was diluted in BSA buffer and injected into the flowcell for tethering, excess DNA was removed with buffer. Imaging buffer consisted of BSA buffer supplemented with 100 mM NaCl, 5 mM MgCl2, 2 mM DTT and gloxy solution (500 units of catalase, 70 units of glucose oxidase and 1% glucose w/v). YOYO-1 was added to the imaging buffer when DNA staining was required.

Total internal reflection fluorescence images were acquired with an inverted Nikon Ti-E microscope. Excitation (488 nM) and emission signals were split with a 638 nm dichroic beam splitter (Chroma) and captured by two EM-CCD cameras (Andor iXon DU897). Images were processed with FIJI software. dCas12a or dCas12a_(FKBP)_ ribonucleoprotein complexes were prepared by mixing the protein with crRNA pools at 1:10 molar ratio in reaction buffer (20 mM Tris-HCl pH 8.0, 100 mM NaCl, 5 mM MgCh, 2% glycerol, 2 mM DTT) followed by incubation at 37°C for 30 min. The complexes were diluted to 10 nM in imaging buffer, injected in the flowcell and incubated at room temperature for 30 min.

Labeling of dCas12a and nucleosomes was carried out *in situ* by injection into the flowcell of anti-FLAG or anti-HA coupled quantum dots (QD_705_), respectively. C-S_10_-B and C-S_10(FRB)_ were labeled via maleimide reaction with Alexa488 and Atto647N, respectively. Both C-S_10_-B and C-S_10(FRB)_ contain an N-terminal cysteine residue and C-S_10(FRB)_ has an additional cysteine in the FRB moiety.

### Dynamic light scattering

Dimerization of dCas12a_(FKBP)_ and C-S_10(FRB)_ via rapamycin was assessed by dynamic light scattering with a Zetasizer μV (Malvern). Both proteins were diluted to 2 μΜ in reaction buffer (20 mM Tris-HCl pH 8.0, 100 mM NaCl, 5 mM MgCl_2_, 2% glycerol, 1 mM DTT) for a total volume of 20 μL and loaded in the cell. Measurements were taken before and immediately after (∼30 s) addition of 5 μΜ rapamycin.

### Electrophoretic mobility shift assay

The crRNA-loaded dCas12a-S_10_ complex was incubated with 100 ng DNA (linear or plasmid) for 1 h before C-S_10_-B addition at stoichiometry ratios ([C-S_10_-B]:[DNA binding sites], N/P) from 0 to 3. Encapsidation was assessed on agarose gel electrophoresis at 4°C and DNA bands were stained with SYBR Green.

### Statistics

All results are given as mean ± standard deviation except for Fig. 1e (mean ± 95% confidence intervals). Non-parametric statistical tests for two (Mann-Whitney) or multiple (Kruskal-Wallis) means were performed on GraphPad Prism. Errors bars for binding distributions of dCas12a, nucleosomes and dCas12a-S_10_ were generated by bootstrap analysis.

## Author Contributions and Notes

A.H.G. and I.J.F. conceived the idea for this study and designed the experiments together with C.C.C. C.C.C. performed all the experiments and analyzed the data jointly with I.J.F. and A.H.G. I.J.F., A.H.G. and C.C.C. wrote the manuscript, which was read and approved by all authors.

This article contains supporting information online.

## Acknowledgments

We thank the CONACyT-University of Texas (CONTEX) grant to carry out this research. A.H.G. also thanks the IA200119 DGAPA-PAPIIT grant. This work was partially funded by the NIH (GM124141 to I.J.F.) and the Welch Foundation (F-1016 to I.J.F.). C.C.C. acknowledges the support of CONACyT for supporting his graduate studies at UT-Austin. We thank David Moreno Gutierrez for providing some of the C-S_10_-B protein and all the members of A.H.G. and I.J.F. labs for valuable discussions. We also thank technicians and facilities of the Institute of Chemistry at UNAM and Tom Wandless at Stanford University for the kind donation of FKBP and FRB plasmids. The authors declare no competing interests.

