## Supplementary figures for "CRISPR-guided programmable self-assembly of artificial virus-like nucleocapsids"

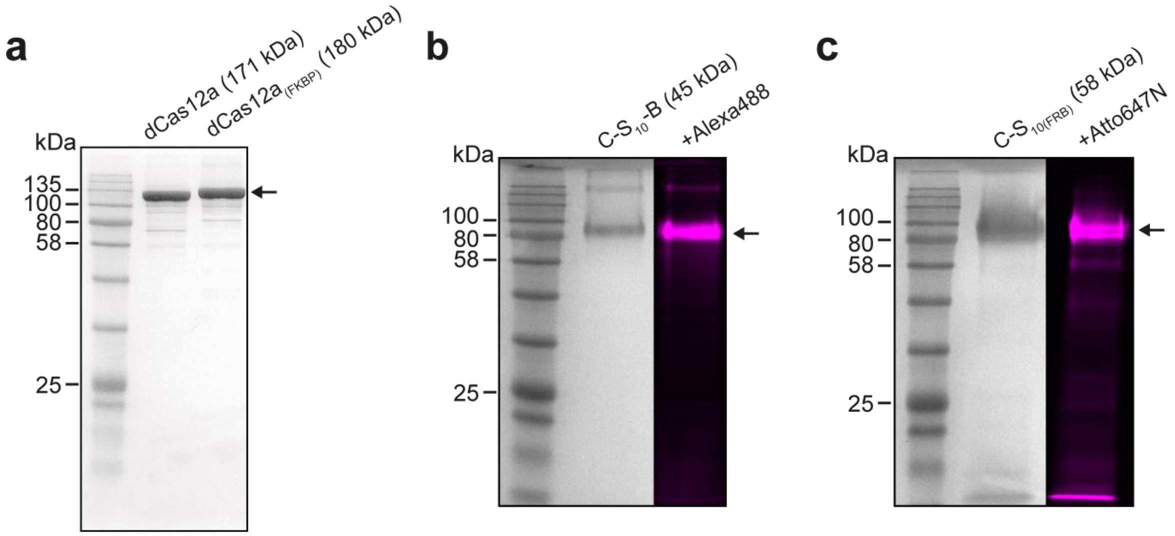


**Figure S1: Proteins used in this study**. **a**: SDS-PAGE gel with dCas12a (171 kDa) and the fusion protein containing dCas12a and the FKBP domain (dCas12a$_{(FKBP)}$, 180 kDa). **b**: SDS-PAGE gel with Coomassie stained (left) and maleimide-Alexa488 labeled (right) C-$S_{10}$-B (45 kDa). **c**: SDS-PAGE gel with Coomassie-stained (left) and maleimide Atto647N labeled (right) fusion protein containing C-$S_{10}$ and the FRB domain (C-$S_{10(FRB)}$) (58 kDa). Both C-$S_{10}$-B and C-$S_{10(FRB)}$ show aberrant migration in gel because of the high proline content (22%) in the C domain.

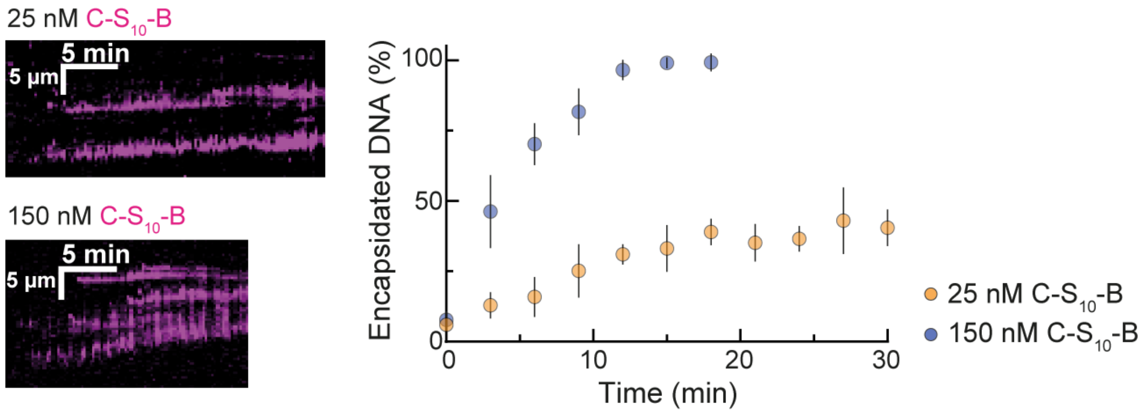


**Figure S2: C-$S_{10}$-B concentration determines nucleocapsid nucleation and growth.** Kymographs showing binding of C-$S_{10}$-B (magenta) on DNA (dark) at 25 nM and 150 nM polypeptide concentration (left) and percentage of the DNA strand length that is encapsidated by the fluorescent C-$S_{10}$-B (right). Shown are the mean and standard deviation for 10 nucleocapsids per condition.

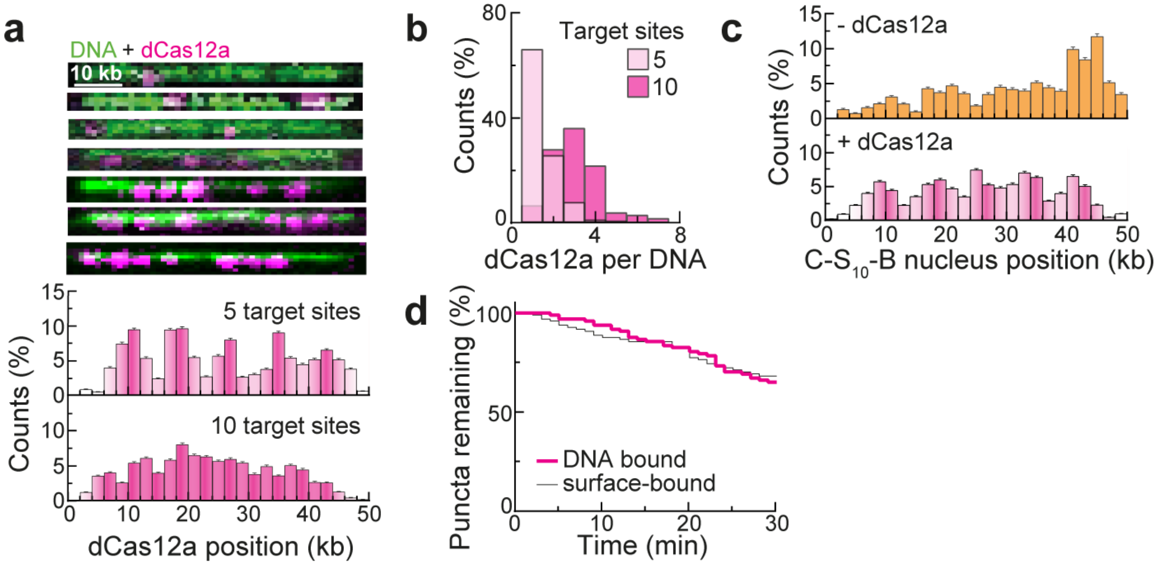


**Figure S3: Decoration of DNA with multiple dCas12a prior to DNA encapsidation by C-$S_{10}$-B. a**: YOYO-1 stained DNA molecules (green) decorated with quantum dot-labeled dCas12a (magenta) (top) and distribution of dCas12a along the DNA (bottom) when binding was targeted to 5 (N = 561 dCas12a molecules) or 10 sites (N = 473 dCas12a molecules). Binding sites are shown with stronger magenta in the histograms. **b**: Number of quantum dot-labeled dCas12a proteins per DNA strand after targeting 5 (N = 561) or 10 (N = 473) binding sites. **c**: Distribution of C-$S_{10}$-B clusters along the DNA with (N = 484) or without (N = 246) decoration with 5 dCas12a. dCas12a binding sites are shown with stronger magenta in the histograms. **d**: Fluorescent quantum dots remaining in the field of view, either bound on the DNA (via dCas12a) or on the lipid surface (N = 100 each) throughout encapsidation with 25 nM C-$S_{10}$-B.

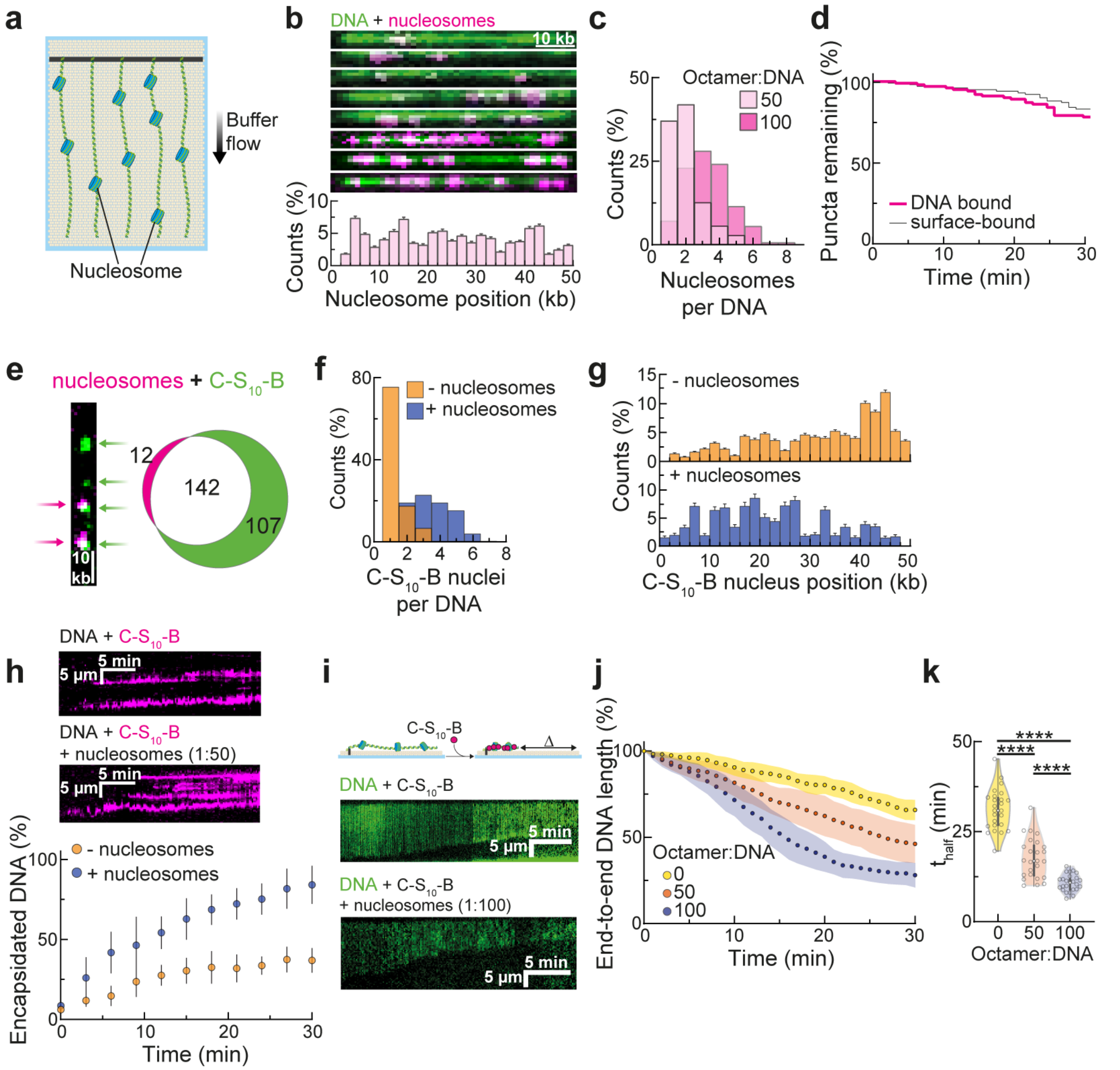


**Figure S4: Prior decoration with nucleosomes improves DNA encapsidation by C-$S_{10}$-B. a**: Human histone octamers were incubated with the DNA at molar ratios 50:1 and 100:1 prior to tethering in the flowcell. **b**: YOYO-1 stained DNA molecules (green) decorated with quantum dot-labeled nucleosomes (magenta) (top) and distribution of nucleosomes on the DNA (bottom) at the 50:1 ratio (N = 279). **c**: Number of quantum dot-labeled nucleosomes per DNA strand at the 50:1 (N = 279) and 100:1 (N = 832) molar ratios. **d**: Fluorescent quantum dots remaining in the field of view, either bound on the DNA (via nucleosomes) or on the lipid surface (N = 100 each) throughout encapsidation with 25 nM

C-$S_{10}$-B. **e**: Double labeling experiment showing co-localization of QD-tagged nucleosomes (magenta, 92%) with fluorescent C-$S_{10}$-B clusters (green, 57%). **f**: Number of C-$S_{10}$-B clusters on naked DNA (N = 246 clusters) versus DNA decorated with nucleosomes at the 50:1 molar ratio (N = 362 clusters). **g**: Distribution of C-$S_{10}$-B clusters along the DNA with (N = 362) or without (N = 246) decoration with nucleosomes at the 50:1 molar ratio. **h**: Kymographs showing C-$S_{10}$-B (magenta) binding on naked or nucleosomes-decorated DNA and percentage of the DNA strand length that is encapsidated by the fluorescent C-$S_{10}$-B. Shown are the mean and standard deviation for 10 nucleocapsids per condition. **i**: Representative kymographs showing faster encapsidation by C-$S_{10}$-B after the DNA (green) is decorated with nucleosomes. **j**: Condensation profiles at 25 nM C-$S_{10}$-B for naked DNA, and DNA decorated with nucleosomes at the 50:1 and 100:1 molar ratios. Shown are the mean and standard deviation for 25 DNA molecules per condition. **k**: Violin plots showing the time ($t_{half}$) required to reach half of maximum condensation for each DNA strand analyzed in (j). Sigmoid curve fitting and extrapolation was used to estimate $t_{half}$ for molecules that did not reach 65.5% encapsidation during the experiment (30 min).

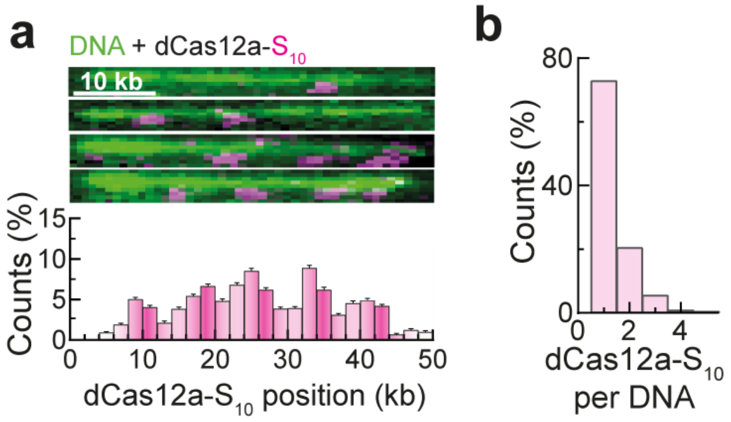


**Figure S5: Decoration of DNA with the previously dimerized complex dCas12a-$S_{10}$. a**: YOYO-1 stained DNA molecules (green) decorated with Atto647N-labeled dCas12a-$S_{10}$ (magenta) (top) and distribution of dCas12a-$S_{10}$ along the DNA (bottom) when binding was targeted to 5 sites (N = 397 dCas12a-$S_{10}$ molecules) along the DNA. Binding sites are shown with stronger magenta in the histograms. **b**: Number of Atto647N-labeled dCas12a-$S_{10}$ per DNA strand after targeting 5 sites (N = 397 dCas12a-$S_{10}$ molecules) along the DNA.

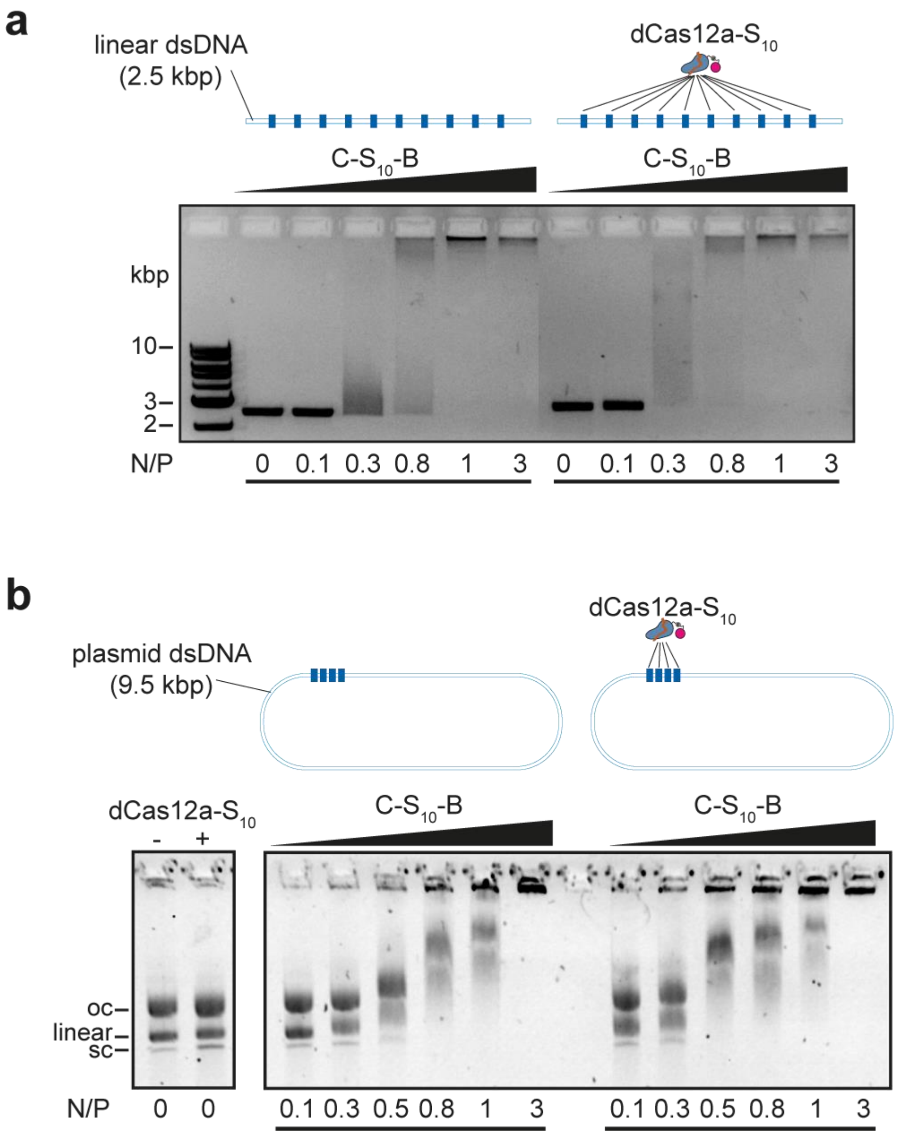


**Figure S6: Positioning multiple dCas12a-$S_{10}$ complexes on DNA substrates improves encapsidation by C-$S_{10}$-B.** C-$S_{10}$-B binding is assessed via electrophoretic mobility shift assays. **a**: A linear 2.5 kbp dsDNA fragment was decorated with ten dCas12a-$S_{10}$ uniformly distributed along the template. Incubation time with C-$S_{10}$-B was 3 h. **b**: dCas12a-$S_{10}$ was directed to four sites on a 9.5 kbp pPIC9 plasmid (in supercoiled, linear and nicked open-circular conformations). Incubation time with C-$S_{10}$-B was 15 h. N/P stands for the stoichiometric ratio between C-$S_{10}$-B and available DNA binding sites (6 bp).

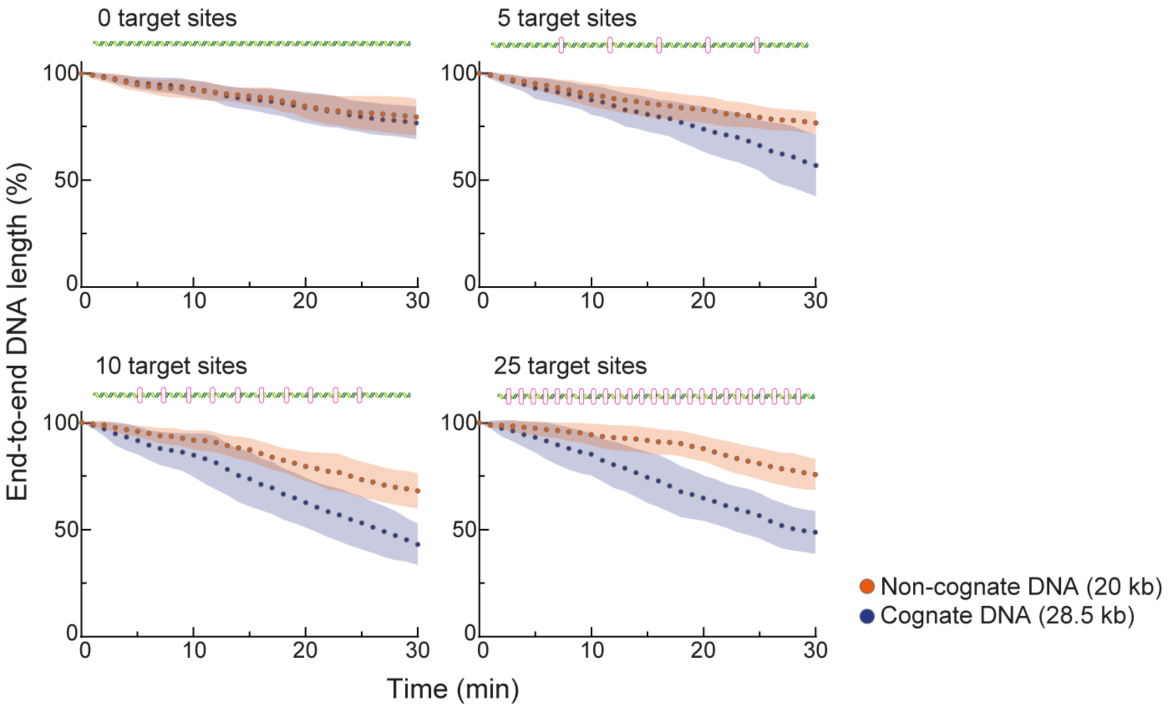


**Figure S7: Encapsidation of cognate and non-cognate DNA templates after selective DNA decoration with dCas12a-S$_{10}$.** Condensation profiles at 10 nM C-S$_{10}$-B for the non-cognate (20 kbp) and cognate (28.5 kbp) DNA strands after dCas12a-S$_{10}$ binding at 0, 5, 10 or 25 sites along the cognate DNA. Circles and shaded areas are the mean and standard deviation for 25 DNA molecules per condition.

**Table S1: Variable sequence of crRNAs used for positioning dCas12a/dCas12a-S10 on the DNA.** All crRNAs consist of a 5’ constant region (UAAUUUCUACUCUUGUAGAU, 20 nt) followed by the 3’ variable region (24 nt) shown in the table.

| | |
|---|---|
| **Full λ DNA** | |
| 5 sites | AUGAUGUUCUGCUGGAUAUGCACU |
| | CCUGACACCGGACGGAAAGCUGAC |
| | AAUGUCGGCUAAUCGAUUUGGCCA |
| | GCUAGCAAUUAAUGUGCAUCGAUU |
| | AUGAACGCAAUAUUCACAAGCAAU |
| 10 sites | CGUGAGAGCUAUCCCUUCACCACG |
| | AUGAUGUUCUGCUGGAUAUGCACU |
| | CGUAUGUCGCCGGAAGACUGGCUG |
| | CCUGACACCGGACGGAAAGCUGAC |
| | UGAUAUGCCGCAGAAACGUUGUAU |
| | AAUGUCGGCUAAUCGAUUUGGCCA |
| | AUGUUCAUCGUUCCUUAAAGACGC |
| | GCUAGCAAUUAAUGUGCAUCGAUU |
| | CCGGACAGGAGCGUAAUGUGGCAG |
| | AUGAACGCAAUAUUCACAAGCAAU |
| **28 kb cognate strand** | |
| 5 sites | UGGCCAAAUCGAUUAGCCGACAUU |
| | GCGUCUUUAAGGAACGAUGAACAU |
| | AAUCGAUGCACAUUAAUUGCUAGC |
| | CUGCCACAUUACGCUCCUGUCCGG |
| | AUUGCUUGUGAAUAUUGCGUUCAU |
| 10 sites | UGGCCAAAUCGAUUAGCCGACAUU |
| | GUCAGAGGCUUGUGUUUGUGUCCU |
| | GCGUCUUUAAGGAACGAUGAACAU |
| | ACUGCGCAUCGCUGGCAUCACCUU |
| | AAUCGAUGCACAUUAAUUGCUAGC |
| | GCUUAAUGACAUUCCUUUCCCGAU |
| | CUGCCACAUUACGCUCCUGUCCGG |
| | CAGGAACGCAACCGCAGCUUAGAC |
| | AUUGCUUGUGAAUAUUGCGUUCAU |
| | AUACCGGAAGCAGAACCGGAUCAC |
| 25 sites | UUGCUUCUCUUGACCGUAGGACUU |
| | CAGUAUUAUGUAGUCUGUUUUUUA |
| | UAAACUCCUUGCAAUGUAUGUCGU |
| | UGGCCUCGAAACCACCGAGCCGGA |
| | CAUCAUUCCAGUCGAACUCACACA |

| | |
|---|---|
| | UCACCGCAGAUGGUUAUCUGUAUG |
| | UCUGGCGAUUGAAGGGCUAAAUUC |
| | CGGGGUGGAUCUAUGAAAAACAUC |
| | AGAAGGAAGAUAUCCUCGCAUGGU |
| | CCAACCAAAUGUAUAUCGAUACCG |
| | ACCCUCAGAGAGAGGCUGAUCACU |
| | ACUUAAUAGUAUUGGUUGCGUAAC |
| | UGAUAUGCCGCAGAAACGUUGUAU |
| | AAUGUCGGCUAAUCGAUUUGGCCA |
| | AUGUUCAUCGUUCCUUAAAGACGC |
| | GCUAGCAAUUAAUGUGCAUCGAUU |
| | CCGGACAGGAGCGUAAUGUGGCAG |
| | AUGAACGCAAUAUUCACAAGCAAU |
| | AGGCCACCGCAUCUCGUGCUGAAG |
| | UUGAAGCAAAUCUGAAACCUAUUA |
| | AGGACACAAACACAAGCCUCUGAC |
| | AAGGUGAUGCCAGCGAUGCGCAGU |
| | AUCGGGAAAGGAAUGUCAUUAAGC |
| | GUCUAAGCUGCGGUUGCGUUCCUG |
| | GUGAUCCGGUUCUGCUUCCGGUAU |
| **Electrophoretic mobility shift assay** | |
| Linear dsDNA (10 sites) | UAGAGCAUAAGCAGCGCAACACCC |
| | AUGAUGAUAUUGAACAGGAAGGCU |
| | UCUCUGCGAGCAUAAUGCCUGCGU |
| | GCCUCCCACGUCUCACCGAGCGUG |
| | UUGAUGGCCUCAUCCACACGCAGC |
| | UAACCGCUUCACACUGACGCCGGA |
| | UUGUUGGUUGCUGCACCAUCCUCU |
| | UGUAUGAAAACGCCCACCAUUCCC |
| | CGGCUCAGUCAUCGCCCAAGCUGG |
| | CGGACACAGUUCCGGAUGGUCAGC |
| Plasmid DNA (4 repeats of 1 site) | CCUUGUUCACCUGGUUGACCAGGG |
